# Phosphate and acidosis cause fiber-type specific changes to cellular and molecular contractile mechanics at 37°C in skeletal muscle from older adults

**DOI:** 10.1101/2025.08.28.672942

**Authors:** Brent A. Momb, Jane A. Kent, Stuart R. Chipkin, Mark S. Miller

## Abstract

Intracellular accumulation of hydrogen ions (H^+^) and inorganic phosphate (P_i_) have temperature-dependent effects on single fiber contractile function between 10-30°C. *In vivo*, human skeletal muscle temperatures range between 35-38°C, and although contractile function is highly dependent on temperature, the effects of fatigue-inducing [H^+^] and [P_i_] on contractile mechanics at 37°C is unknown. Using sinusoidal analysis, the independent and combined effects of these metabolites on cellular and molecular contractile function were determined at 37°C in slow-contracting myosin heavy chain (MHC) I and fast-contracting MHC IIA fibers from vastus lateralis muscle of 13 older adults (8 females), under four conditions: maximal calcium activation (“control”; 5 mM P_i_, pH 7.0), high P_i_ (30 mM), low pH (6.2), and fatigue (30 mM P_i_ and pH 6.2). Specific tension (force/cross-sectional area, mN/mm^2^) in both fiber types was reduced only under fatigue conditions (20-26%). MHC I fibers had slower cross-bridge kinetics with fewer or less stiff strongly-bound myosin-actin cross-bridges in high P_i_, low pH, and fatigue. In contrast, fatigued MHC IIA fibers had faster cross-bridge kinetics with increased myofilament and/or cross-bridge viscosity. Single fiber oscillatory work was reduced in both fiber types when P_i_ or pH alone was altered. However, fatigue conditions returned oscillatory work values toward control through alterations to cross-bridge kinetics in MHC I fibers and changes to work absorption and production processes in MHC IIA fibers. These findings quantify fiber-type specific mechanical and kinetic mechanisms of fatigue in human skeletal muscle at 37°C, thus advancing our understanding of metabolite-based muscle fatigue *in vivo*.

**KEY POINTS SUMMARY:**

- Working skeletal muscle increases intracellular concentrations of hydrogen ion and inorganic phosphate, leading to fatigue, or loss of force-generating capacity
- Temperature plays a well-established role in the muscle response to hydrogen ion and/or inorganic phosphate accumulation, but has not previously been studied at human body temperature (37°C)
- At 37°C, reduced force generation only occurs when high phosphate and hydrogen ions are combined, not when changed individually
- In slow-contracting fibers, fatigue slowed myosin-actin cross-bridge kinetics and reduced the number or stiffness of strongly-bound cross-bridges. In fast-contracting fibers, fatigue increased myosin-actin cross-bridge kinetics and increased myofilament viscosity.
- The distinct responses by fiber type to fatigue provides new insight into its mechanisms and advances our understanding of the whole muscle and body responses to fatigue

**Abstract Figure:** 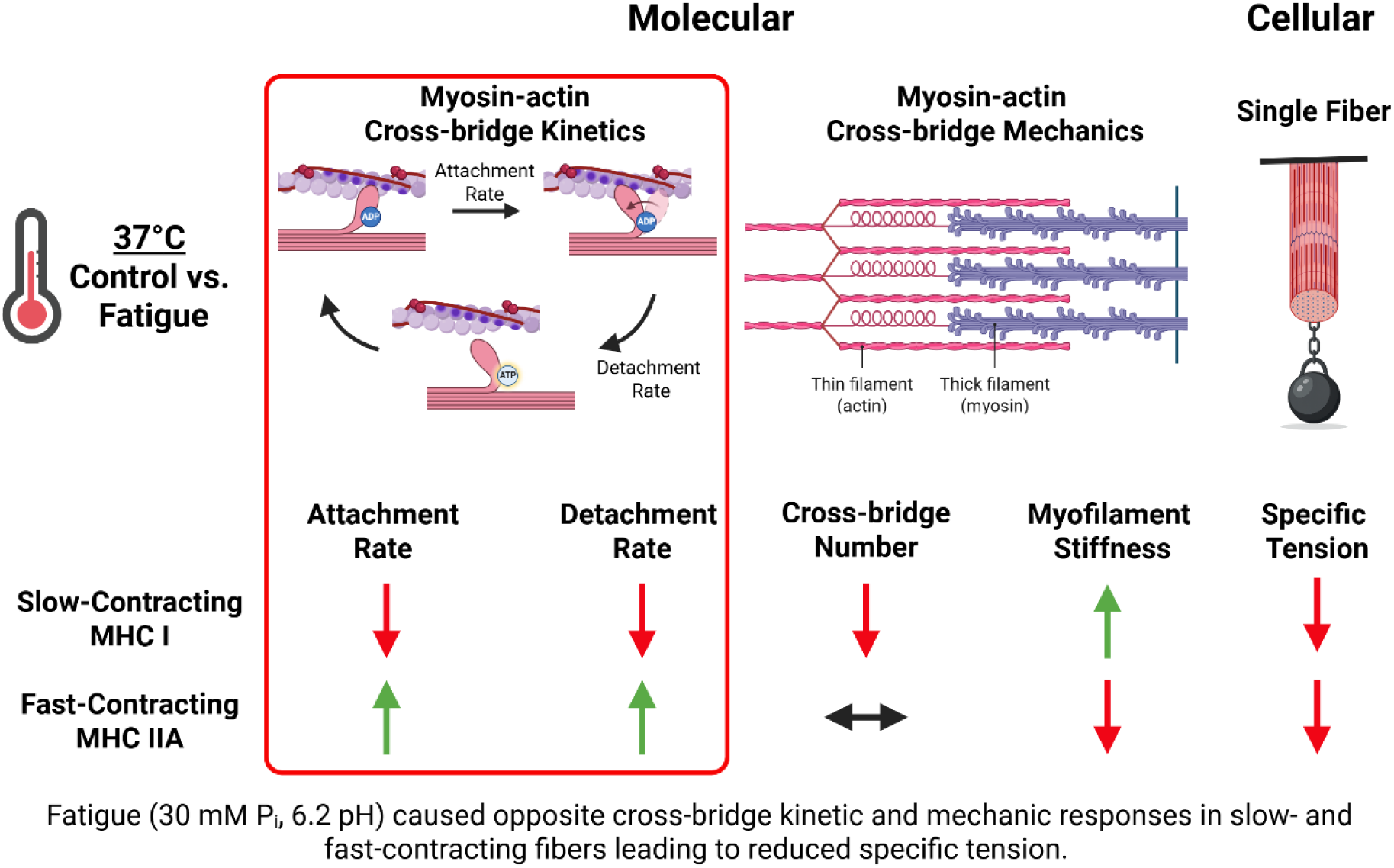

## INTRODUCTION

In working human skeletal muscle, contractile failure can occur due to the intracellular accumulation of inorganic phosphate (P_i_) and acidosis (high H^+^ or reduced pH) (Cady *et al*., 1989; Weiner *et al*., 1990; Kent-Braun, 1999; Broxterman *et al*., 2017; Fitzgerald *et al*., 2023), which directly alter muscle contractile mechanics at the cellular and molecular levels (Sundberget al., 2018, 2025; Foster et al., 2025) and thus, result in a loss of force-generating capacity, or fatigue (Fitzgerald *et al*., 2023). Although both P_i_ and pH have temperature-dependent effects on contractile function, the impact of these metabolites in single fibers has primarily been studied between 10-30°C in animal models (Cooke *et al*., 1988; Coupland *et al*., 2001; Debold *et al*., 2004; Knuth *et al*., 2006; Karatzaferi *et al*., 2008). *In vivo*, human skeletal muscle temperatures range between 35-39°C (Bergh & Ekblom, 1979; Kenny *et al*., 2003; Flouris *et al*., 2015). Further, the fatigability of different myosin heavy chain (MHC) isoforms has been of great interest in the field of muscle physiology as the proportion of slow-or fast-contracting fibers has been linked to the development of whole-muscle contractile failure (Thorstensson & Karlsson, 1976). Despite this, potential fiber-type specific responses to fatigue have not been studied extensively. Since the accumulation of P_i_ and reduced pH play a large role in muscle fatigue (Kent-Braun *et al*., 1993, 2002; Cooke, 2007; Broxterman *et al*., 2017; Vandenboom, 2017), have temperature-sensitive effects on contractile function (Nelson *et al*., 2014; Sundberg *et al*., 2018), and may cause different responses depending on fiber type (Mizuno *et al*., 1994*b*, 1994*a*), understanding how myosin and actin function at 37°C is crucial to understanding muscle fatigue.

The combined effects of high P_i_ and low pH on contractile function has been documented in single fibers from animals (Cooke *et al*., 1988; Potma *et al*., 1995; Nocella *et al*., 2011; Nelson *et al*., 2014) and humans (Sundberg et al., 2018, 2025; Foster et al., 2025), with studies indicating that their effect on force or contractile velocity generally diminishes as temperature is increased toward a physiological range 15-30°C (Nelson *et al*., 2014; Sundberg *et al*., 2018).

Lower force in single fibers during fatigue induced by high P_i_ and low pH has been thought to be due to reduced force per cross-bridge (Nocella *et al*., 2011; Nelson *et al*., 2014; Foster *et al*., 2025) and/or reductions in cross-bridge number (Nocella *et al*., 2011; Foster *et al*., 2025), and may be fiber-type specific (Foster *et al*., 2025). While not all studies agree on the underlying mechanism, slowed velocity is generally thought to be due to slowed cross-bridge kinetics (Nelson *et al*., 2014; Sundberg *et al*., 2018; Foster *et al*., 2025). Until recently, only one research group had performed experiments on single fibers in humans at 30°C (Sundberg *et al*., 2018), and solely in slow-contracting MHC I fibers, which may respond to fatigue conditions differently than MHC IIA (Potma *et al*., 1995; Foster *et al*., 2025). Thus, a lack of information exists as to how the combination of P_i_ and pH impact cellular- and molecular-level mechanics in MHC I and MHC IIA fibers at *in vivo* temperature.

While P_i_ and pH generally do not change independently *in vivo,* the contractile response to each metabolite individually may provide insight into their combined behavior by understanding which steps in the cross-bridge cycle are altered. The effects of P_i_ or pH alone on contractile function has been examined extensively in single fibers from animals (Cooke & Pate, 1985; Metzger & Moss, 1987, 1990; Zhao & Kawai, 1994; Wahr *et al*., 1997; Westerblad *et al*., 1997; Wang & Kawai, 2001; Widrick, 2002; Debold *et al*., 2004; Knuth *et al*., 2006; Karatzaferi *et al*., 2008). However, limited data exist on metabolites alone in humans as only one study has examined the effect of P_i_ alone and only in a small concentration range (0-4 mM P_i_) (Sundberg *et al*., 2025), although P_i_ has been shown to increase to 30 mM or more *in vivo* (Cady et al., 1989; Weiner et al., 1990; Kent-Braun, 1999; Broxterman et al., 2017; Fitzgerald et al., 2023).

Obtaining data in humans is important as humans and animals have different skeletal muscle contractile properties (Pellegrino *et al*., 2003), including myosin-actin cross-bridge kinetics, which tend to slow as species increase in size (Nyitrai *et al*., 2006). Additionally, P_i_ (Coupland *et al*., 2001; Debold *et al*., 2004) and pH (Pate *et al*., 1995; Wiseman *et al*., 1996; Westerblad *et al*., 1997) have temperature-dependent effects, such that as temperature increases their independent effects on single fiber force production and contractile velocity are diminished.

Furthermore, rates of myosin attachment and detachment are highly temperature-sensitive (Zhao & Kawai, 1994; Wang & Kawai, 2001), and directly impacted by the accumulation of P_i_ or H^+^ (Zhao & Kawai, 1994; Wang & Kawai, 2001; Debold *et al*., 2008; Woodward & Debold, 2018; Foster *et al*., 2025; Marang *et al*., 2025). Thus, studying these metabolites at 37°C in humans will give insight into their respective roles in muscle fatigue under conditions that mimic those *in vivo*.

To date, the study of human skeletal muscle tissue at physiological temperatures has been limited to some extent due to deterioration of MHC IIA fibers above 15-25°C and MHC I fibers above 30°C (Sundberg *et al*., 2018). To achieve a higher testing temperature of 37°C and maintain fiber stability in human single fibers, we chemically fix the ends of single fibers with a glutaraldehyde solution to cross-link proteins (Miller *et al*., 2010). To quantify cross-bridge mechanics and kinetics, we use small-amplitude (0.05% of fiber length) sinusoidal perturbations of single fibers to reduce damage to fibers. Fiber damage can occur with other techniques that require large length changes (Sundberg *et al*., 2018), such as force-velocity curves or unloaded shortening velocity measurements. Importantly, the sinusoidal analysis method permits the assessment of myofilament protein function in its native, three-dimensional configuration at the level of the myosin-actin cross-bridge. Under calcium-activated conditions, sinusoidal length perturbations, applied below the unitary myosin step size and across a range of frequencies (Kawai, 2003; Brenner, 2006), facilitate the measurement of muscle mechanical properties relevant to specific steps in the cross-bridge cycle (Zhao & Kawai, 1993; Kawai *et al*., 1993; Mulieri *et al*., 2002; Palmer *et al*., 2007). The application of sinusoidal analysis at 37°C with fixation of single fibers in human skeletal muscle thus allows for a mechanistic understanding of the impact of elevated P_i_ and reduced pH on muscle contractile function under more physiologically relevant conditions.

The purpose of this study was to quantify the independent and combined effects of elevated P_i_ and reduced pH on cellular (single fiber) and molecular (myosin-actin cross-bridge kinetics and mechanics) contractile function at 37°C in single human skeletal muscle fibers from healthy older adults. Slow-contracting myosin heavy chain (MHC) I and fast-contracting MHC IIA fibers from the vastus lateralis muscle were examined under four conditions of maximal calcium activation: control (5 mM P_i_, pH 7.0), high P_i_ (30 mM), low pH (6.2), and fatigue (30 mM P_i_ and pH 6.2). Control phosphate (P_i_) concentration was 5 mM to align with resting [P_i_] in healthy human gastrocnemius (5 mM) and quadriceps (4.5 mM) muscles (Pathare *et al*., 2005; Kemp *et al*., 2007). Fatigue values for P_i_ and pH were chosen as P_i_ has been shown to increase to 30 mM or more and pH to decline to as low as 6.2 *in vivo* during fatiguing contraction protocols in various human skeletal muscles expressing a range of MHC isoforms (Cady *et al*., 1989; Weiner *et al*., 1990; Kent-Braun, 1999; Broxterman *et al*., 2017; Fitzgerald *et al*., 2023). Skeletal muscle biopsies were performed in healthy older (65-80 years) participants with habitual physical activity was measured by accelerometry.

## MATERIALS AND METHODS

### Ethical Approval

Written, informed consent was obtained from each volunteer before their participation. The protocol was approved by the Institutional Review Board at the University of Massachusetts Amherst and was performed following the *Declaration of Helsinki*.

### Participants

Thirteen healthy older adults (5 males, 8 females) completed this study. All volunteers were aged 65-80 years, generally healthy by self-report, ambulatory without the use of walking aids, living independently in the community, and sedentary to somewhat active (no more than two 30-minute light- to moderate-volitional exercise sessions per week by self-report) as verified by accelerometry. Females were postmenopausal (cessation of menses > 1 year). Individuals were excluded if they had a history of major neurological or neuromuscular conditions that could impact physical function, history of myocardial infarction, angina, peripheral vascular disease, surgical or percutaneous coronary artery revascularization, history of severe pulmonary disease, history of rheumatoid arthritis, history of diabetes or other metabolic disease that could impact neuromuscular function, uncontrolled hypertension (blood pressure > 140/90 mmHg), smoking in the past year, moderate to severe lower extremity arthritis, pain, muscle cramps, joint stiffness, light-headedness or other symptoms upon exertion, the use of beta-blockers, sedatives, tranquilizers, or other medication that could impair physical function, body mass index >35 kg/m^2^ as increased fat mass may alter single muscle fiber performance, or body mass index <18 kg/m^2^. Any persons taking anti-coagulant medication or with known coagulopathies due to increased bleeding risk from biopsy procedure, participants with a contraindication for magnetic resonance testing including a pacemaker or other implant, males and females undergoing hormone replacement therapy because this treatment may circumvent normal age-related declines in sex hormone levels (if taken, hormone therapy must have been > 5 years ago), unintentional weight loss greater than 2.5 kg during the last 3 months, currently participating or have participated in a weight loss or exercise training program in the last year, inability to understand written and spoken English, and inability to follow instructions as determined by investigators during the consenting process, were also reasons for exclusion. All participants obtained clearance from their health care provider to participate. Eligibility was determined during a screening visit, followed by collection of informed consent and anthropometric data.

### Accelerometry

To account for the potential confounder of physical activity on skeletal muscle function, participants were asked to complete a physical activity log and wear an accelerometer (ActiGraph GT3X-BT) on their dominant hip for 7 days. Data were collected at 60 Hz in 1 second epochs. ActiLife software (ActiGraph, Pensacola, FL) was used to calculate moderate-to-vigorous physical activity (MVPA) using the Troiano cut points (Troiano *et al*., 2008) from a minimum of four days, including one weekend day. All days included at least 10 hours of wear time.

### Experimental Solutions

All solutions used for biopsy processing and single fiber experiments were calculated using the equations and stability constants according to Godt and Lindley (Godt & Lindley, 1982). Characteristics of the solutions used for dissection, skinning, storage, and single fiber experiments are given in Table 1.

**Table 1.**
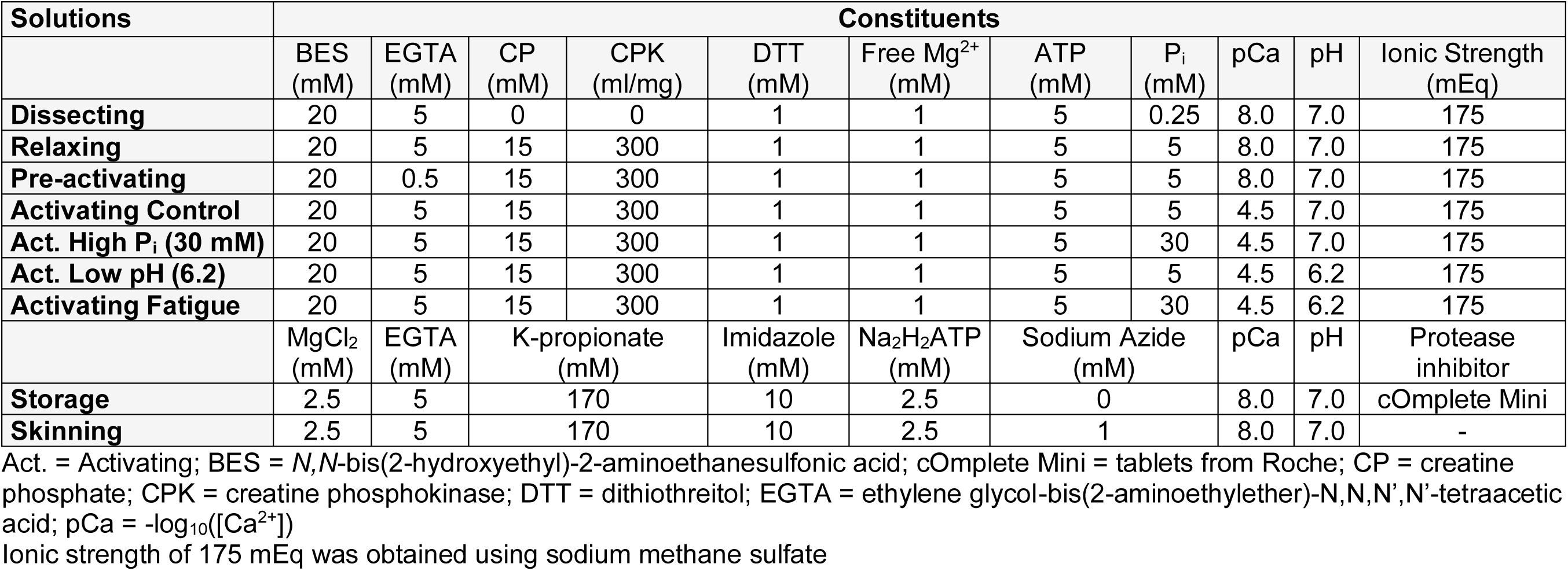
Characteristics of solutions used for skeletal muscle fibers.

### Skeletal Muscle Tissue Biopsy and Processing

Volunteers were fasted for at least 10 h prior to the muscle biopsy. Muscle tissue from one or two legs was obtained via percutaneous biopsy of the vastus lateralis under lidocaine anesthesia. Skeletal muscle tissue was placed immediately into cold (4°C) dissecting solution. Muscle bundles were placed in skinning solution for 24 h at 4°C and were then placed in storage solution with increasing concentration of glycerol (10% v/v glycerol for 2 h, 25% v/v glycerol for 2 h) until reaching the final storage solution (50% v/v glycerol) and incubated at 4°C for 18–20 h. Thereafter, bundles were stored at −20°C until isolation of single fibers for mechanical measurements, which occurred within 4 weeks of the biopsy.

### Preparation of Single Fibers for Mechanical Analysis

Single fiber preparation has been described previously (Miller *et al*., 2010). Briefly, muscle bundles were incubated in dissection solution containing 1% Triton X-100 (v/v) for 30 min at 4°C. Segments (∼2.0-2.5 mm) of single fibers were manually isolated from muscle bundles, with aluminum T-clips placed at both ends of the fiber. The fiber was again incubated in dissecting solution containing 1% Triton X-100 (v/v) for 30 min at 4°C to ensure removal of the sarcolemma and sarcoplasmic reticulum. The fiber was then mounted onto hooks in dissecting solution at room temperature on the CSA/fixation rig. Top and side diameter measurements were made at three positions along the length of the middle 1-2 mm of the fiber using a filar eyepiece micrometer (Lasico, Los Angeles, CA, USA) and a right-angled, mirrored prism to obtain the side-to-top width ratio. Fibers were then fixed at two points approximately 1-2 mm apart with glutaraldehyde, at the end of each T-clip. New T-clips were placed on the fixed regions dyed with bromophenol blue. The fiber was then transferred to the sinusoidal analysis rig for mechanical testing.

### Experimental Protocol

Each fiber was tested for maximal specific tension (mN/mm^2^) at 37°C in each of the experimental conditions in a random order with minimal time (< 5 s) between each trial: Control (P_i_ = 5 mM, pH = 7.0), high P_i_ (P_i_ = 30 mM, pH = 7.0), low pH (P_i_ = 5 mM, pH 6.2), fatigue (P_i_ = 30 mM, pH = 6.2). Myofilament mechanical properties, including myosin-actin cross-bridge kinetics, were measured using small amplitude sinusoidal analysis in the same randomized order as the initial measurements for maximal specific tension. Sinusoidal analysis experiments took ∼40 s to complete at 37°C.

### Single Fiber Mechanical Analysis

Single muscle fiber mechanical analysis was adapted from a previous study (Miller *et al*., 2010). The T-clipped ends of the fiber were attached to a piezoelectric motor (Physik Instrumente, Auburn, MA, USA) and a strain gauge (SensorNor, Horten, Norway) on the sinusoidal analysis rig in relaxing solution at 15°C to maintain fiber integrity. The fiber was manually stretched until the sarcomere length was 2.65 μm (IonOptix, Milton, MA, USA), as this resembles *in vivo* sarcomere length in human skeletal muscle (Chen *et al*., 2016). A camera (Point Grey, FLIR Integrated Imaging Solutions, Inc., Richmond, BC, Canada) was used to image the fiber and a computer program (ImageJ) performed a fast-fourier transformation of the visible light and dark sarcomere patterns to determine sarcomere length. Fiber length (L) was measured using a micrometer as the distance between the inside edges of the T-clips on each end of the fiber. Fiber top width was measured in the middle of the fiber and side width was estimated using the side-to-top width ratio obtained on the CSA/fixation rig. Fiber top and estimated side widths were used to calculate elliptical CSA.

Starting in relaxing solution at 15°C, the fiber was slackened completely, the force gauge zeroed, the fiber pulled back to its original position, allowed to equilibrate for 1 min and relaxed isometric force measured (e.g., setting the force baseline). The fiber was transferred to pre-activating solution for 30 s and then to control activating solution, with force recorded at its plateau. The fiber was returned to relaxing solution, given time to relax, and stretched as necessary to return sarcomere length to 2.65 μm. Starting again in relaxing solution, temperature was increased to 37°C over the course of 1-2 min, force baseline was set, the fiber was transferred from pre-activating to one of the activating solutions using the above procedure, and specific tension recorded at its plateau. The fiber was immediately moved to the other activation conditions (control, high P_i_, low pH, or fatigue) in a randomized order to obtain their specific tension values. After each specific tension was recorded, the fiber was transferred to each activating solution in the same randomized order with small-amplitude, sinusoidal length changes (0.05% of L) applied to the fiber at 42 frequencies (3 to 200 Hz). Length and force were normalized to determine fiber strain (ΔL/L) and stress (force/CSA). Elastic (E_e_) and viscous (E_v_) moduli (N/mm^2^) were calculated from the stress transient by determining the magnitudes of the in-phase and out-of-phase components (0° and 90° with respect to strain). The elastic and viscous moduli are the real and imaginary parts of the complex modulus, i.e., the ratio of the stress response to the strain.

Myosin-actin cross-bridge mechanics and kinetics as well as properties of the myofilaments were derived, as previously described (Miller *et al*., 2010; Momb *et al*., 2022), by fitting the complex modulus with the following six-parameter equation:

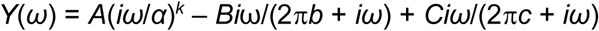

where *ω* = 2π*f* in s^-1^, *A*, *B* and *C* are magnitudes expressed in N/mm^2^, 2π*b* and 2π*c* are characteristic rates expressed in s^-1^, *i* = −1^1/2^, *α* = 1 s^-1^ and *k* is a unitless exponent. This analysis yields three characteristic processes, A, B and C, which relate to various mechanical (*A*, *B*, *C* and *k*) and kinetic (2π*b* and 2π*c*) properties of the crossbridge cycle. Nyquist plots, or viscous versus elastic modulus, show the stress response of the fiber to strain over a range of frequencies, which can be fit using the six-parameter equation above. The A-process (*A* and *k*) has no enzymatic dependence and, under Ca^2+^-activated conditions, reflects the viscoelastic mechanical response of the structural elements of the muscle fiber (the myofilament lattice stiffness and attached myosin heads in series) (Mulieri *et al*., 2002; Palmer *et al*., 2004). The parameter *A* indicates the magnitude of the viscoelastic modulus, whereas *k* represents the degree to which these viscoelastic magnitudes are purely elastic (*k* = 0) or purely viscous (*k* = 1). The B- and C-process magnitudes (*B* and *C*) are proportional to the number of strongly bound myosin-actin interactions and/or the stiffness of the cross-bridges (Kawai *et al*., 1993).

The *C* / *B* ratio is the relationship between the magnitude of the work-absorbing, or positive viscous modulus, C-process (*C*) and the work-producing, or negative viscous modulus, B-process (*B*). The frequency portion of the B-process (2π*b*) is interpreted as the rate of myosin transition from the weakly to strongly bound state, or the (apparent) rate of myosin force production (Zhao & Kawai, 1993). The frequency portion of the C-process (2π*c*) represents the cross-bridge detachment rate (Palmer *et al*., 2007). The 2π*c* / 2π*b* ratio is the ratio between the rate of detachment to the rate of force production. From the sinusoidal analysis, work (J/m^3^) was calculated as work = π *f t* (−*E_v_*) (*L_amp_*)^2^ where *f* is the frequency of the length perturbations (Hz), *t* is the time needed to perform the length perturbations (*s*), *E_v_* is the viscous modulus (N/mm^2^), and the fractional change in length (*L_amp_*) is 0.0005% of fiber length. Of note, work is positive when the viscous modulus plot is negative.

### MHC Isoform Identification

After sinusoidal analysis measurements, single fibers were placed in 30 µl loading buffer, heated for 2 minutes at 65°C, and stored at −80°C until determination of MHC isoform composition by SDS-PAGE to identify fiber type, as described (Miller *et al*., 2010).

### Statistical Analyses

All data are reported as mean ± standard deviation (SD) and differences were considered significant at *p* ≤ 0.05. As there were multiple observations within the same participant (i.e., multiple fibers per participant), main effects for each fiber type were determined using a repeated-measures linear mixed model, with a random effect, to account for grouping observations within participants, as reported previously (Miller *et al*., 2013; Momb *et al*., 2022), and a repeated effect to account for the between-participant factor (Control, high P_i_, low pH, and fatigue). If a main effect was noted, Fisher’s Least-Significant Difference post-hoc test was performed to determine pairwise differences between the experimental conditions. All analyses were conducted using IBM SPSS Statistics for Windows version 25.0 (IBM, Armonk, NY).

## RESULTS

### Participant Characteristics

Thirteen older adults (8 female) aged 70.8 ± 1.9 yr (range 66-76) were included in this study. The participants had a height of 171.4 ± 4.2 cm (range 160-178) body mass of 75.2 ± 4.5 kg (range 51.8-98.7), and body mass index of 25.6 ± 1.4 kg/m^2^ (range 18.3-31.8). Daily step count averaged 6,481 ± 2,342 (range 3,680-11,181), weekly moderate-to-vigorous activity was 282 ± 111 minutes (range 124-523), and daily activity counts of 417 ± 58 arbitrary units (range 350-539).

### Cellular-Level Outcomes

Manipulating P_i_ or pH independently did not change specific tension compared with control conditions in MHC I (P_i_: *p* = 0.0942, pH: *p* = 0.0655) or IIA fibers (P_i_: *p* = 0.413, pH: *p* = 0.260) (Figure 1). In contrast, altering both P_i_ and pH in solution (fatigue) reduced specific tension in MHC I (20%, *p* < 0.001) and IIA (26%, *p* < 0.001) fibers compared with control conditions. These results indicate that an interdependent effect occurs with the combination of high P_i_ and low pH compared to either condition alone, and that this effect is robust at physiological temperature.

**Figure 1.**
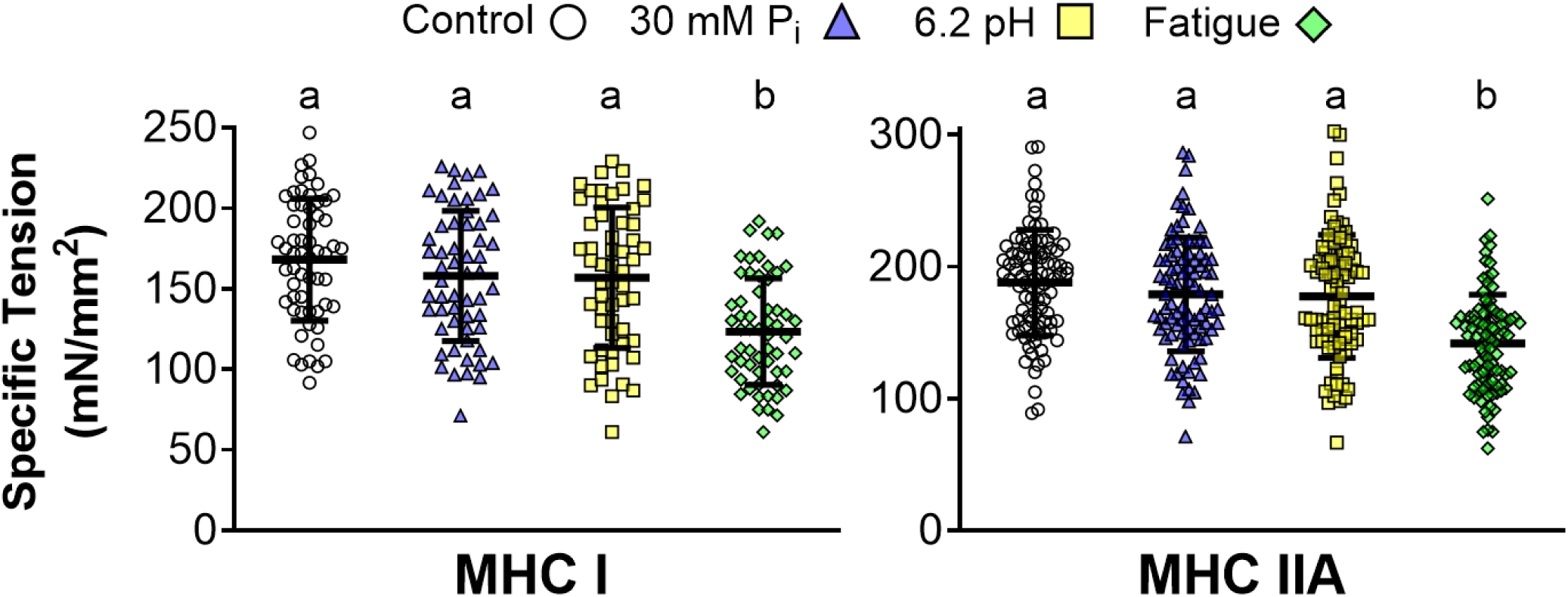
**Single-fiber maximum Ca^2+^-activated (pCa 4.5) specific tension by MHC isoform for Control, high P_i_, low pH, and fatigue conditions at 37°C.** Data are presented as mean ± SD with individual fibers as data points, letters above bars indicate significant (*p* < 0.05) pair-wise differences between conditions for MHC I or IIA.

### Molecular-Level Cross-bridge Mechanical Analysis

To understand the molecular mechanisms of force production, the six-parameter model was used to fit the sinusoidal analysis data. The mechanical analysis, involving parameters *B* and *C*, indicated that the number and/or stiffness of strongly-bound myosin-actin cross-bridges were reduced with greater P_i_ or lower pH in MHC I (all *p* < 0.001) and MHC IIA fibers (all *p* < 0.001) (Figure 2.1-2.2). The myofilament properties analysis showed that parameter *A* increased and *k* decreased under conditions of high P_i_ or low pH in MHC I (A: P_i_: *p* = 0.00258, pH: *p* = 0.00908; *k*: P_i_: *p* = 0.00375, pH: *p* = 0.0413) and IIA fibers (*A*: P_i_ *p* = 0.0182; pH *p* < 0.001; *k*: pH *p* = 0.00378), with the exception that *k* remained unchanged for high P_i_ in MHC IIA (*p* = 0.385) (Figure 2.3-2.4). These alterations to *A* and *k* indicate that P_i_ and pH each independently increase the myofilament stiffness of MHC I and IIA fibers.

**Figure 2.**
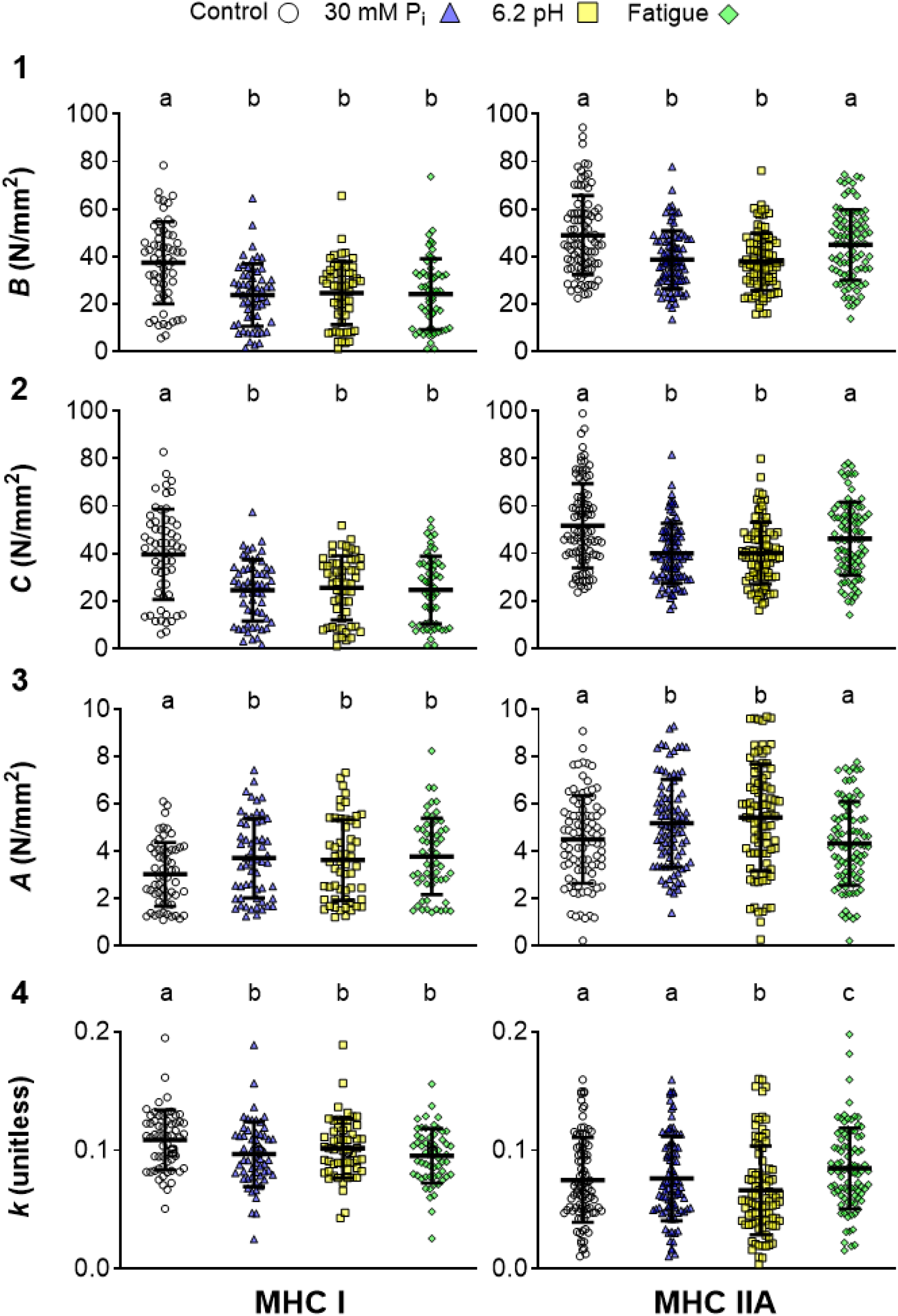
**Single-fiber maximum Ca^2+^-activated (pCa 4.5) sinusoidal analysis model mechanical parameters by MHC isoform for Control, high P_i_, low pH, and fatigue conditions at 37°C**. (1-2) Parameters *B* and *C* are proportional to the number of myosin heads strongly bound to actin and/or cross-bridge stiffness. (3) *A* and (4) *k* represent the underlying stiffness of the lattice structure and the attached myosin heads in series. Data are presented as mean ± SD with individual fibers as data points. Letters above bars indicates significant (*p* < 0.05) pair-wise differences between conditions for MHC I or IIA.

Under fatigue conditions, MHC I fibers displayed reductions in *B* and C compared to control (*p* < 0.001) with a concomitant increase in *A* (*p* = 0.00117) and decrease in *k* (*p* < 0.001) (Figure 2, left panels). To understand why specific tension dropped in MHC I with fatigue despite the apparently “off-setting” mechanical and myofilament changes described above, the percent change of *B* and *C* from control, or Δ*B* and Δ*C*, was determined for each condition as parameters *B* and *C* inherently have a high level of variability in the measurement. Expression as a percent change shows the consistency and magnitude of change within individual fiber experiments. The fatigue condition produced greater decreases in Δ*B* and Δ*C* (33-35%) (both *p* < 0.001) relative to the independent manipulations of P_i_ and pH (21-27%) (*p* = 0.411-0.519) (Figure 3.1-3.2, left panels). These results indicate the reduction in specific tension with fatigue is due to fewer strongly-bound myosin-actin cross-bridges and/or their stiffness compared with P_i_ or pH alone. The percent change was examined for the other parameters *A* and *k*, but parameters *B* and *C* were the only variables that had significantly different responses in these measures across conditions.

**Figure 3.**
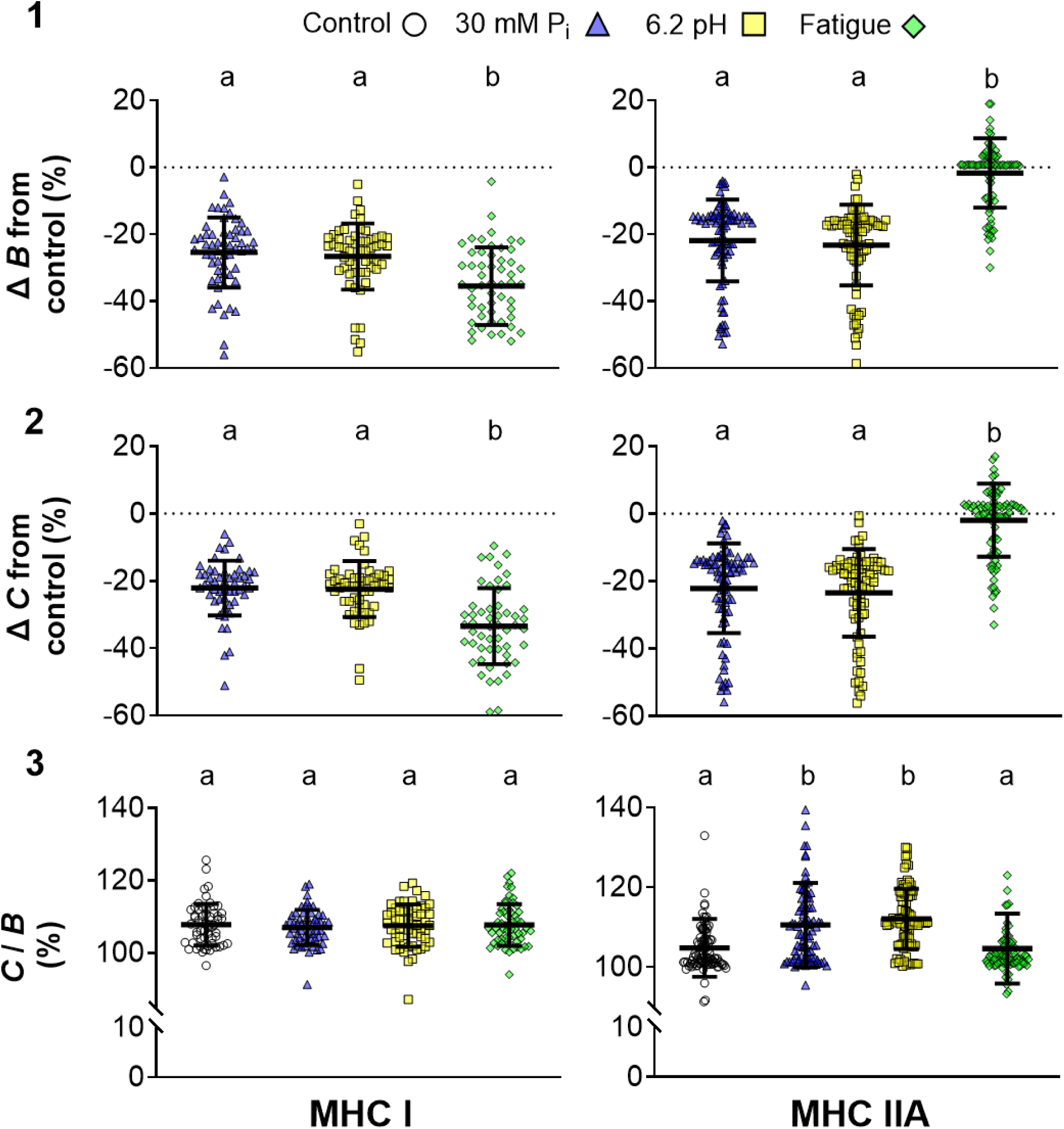
**Relative changes in sinusoidal analysis model parameters *B* and *C* by MHC isoform for Control, high P_i_, low pH, and fatigue conditions at 37°C.** (1-2) Parameters *B* and *C* are proportional to the number of myosin heads strongly bound to actin and cross-bridge stiffness during maximum Ca^2+^ activation, respectively. (3) *C* / *B* is the ratio of parameter C (the magnitude of work absorption) and parameter *B* (the magnitude of work production). Single fiber data points are overlayed on top of bars. Data are presented as mean ± SD with individual fibers as data points, letters above bars indicate significant (*p* < 0.05) pair-wise differences between conditions for MHC I or IIA. Horizontal dotted lines in 1 and 2 indicates zero relative change in *B* or *C*.

In MHC IIA fibers, fatigue conditions increased *k* (*p* < 0.001) with no change in parameters *B* (*p* = 0.285)*, C* (*p* = 0.0834), or *A* (*p* = 0.381) (Figure 2, right panels), suggesting that the drop in specific tension with fatigue was due to fibers becoming more viscous, or work-absorbing. The fatigue condition did not significantly change Δ*B* or Δ*C* relative to control, as expected as there was no change in *B* and *C*, and was higher than the P_i_ and pH conditions (*p* < 0.001). The independent manipulations of P_i_ and pH decreased Δ*B* and Δ*C* to a similar extent (21-24%; *p* = 0.387-0.482), as expected due to their lower *B* and *C* values. The ratio between the magnitude of the work-absorbing, or positive viscous modulus, C-process (*C*) and the work-producing, or negative viscous modulus, B-process (*B*) provides potential insight into fatigue-induced alterations in contractile properties. The *C* / *B* ratio was unchanged in MHC I fibers under any condition (*p* = 0.475-0.944) (Figure 3.3, left panel). In contrast, in MHC IIA fibers this ratio was increased for P_i_ (*p* < 0.001) and pH (*p* < 0.001) alone but not combined (*p* = 0.904) (Figure 3.3, right panel). A greater *C* / *B* ratio indicates that the work-absorbing processes became relatively larger, a result that will become more relevant when examining the changes in oscillatory work production.

### Molecular-Level Cross-bridge Kinetic Analysis

In MHC I fibers, the rates of myosin force production (2π*b*) (P_i_: *p* = 0.0234, pH: *p* < 0.001, fatigue: *p* < 0.001) and cross-bridge detachment (2π*c*) (P_i_: *p* = 0.0453, pH: *p* < 0.001, fatigue: *p* = 0.00848) slowed in all conditions compared with control (Figure 4.1-4.2). The fatigue and low pH conditions had the slowest 2π*b*, whereas 2π*c* was most altered by low pH alone. This latter result suggests that, in fatigue, high P_i_ has a moderating effect on the slowing of cross-bridge detachment due to acidosis in MHC I fibers.

**Figure 4.**
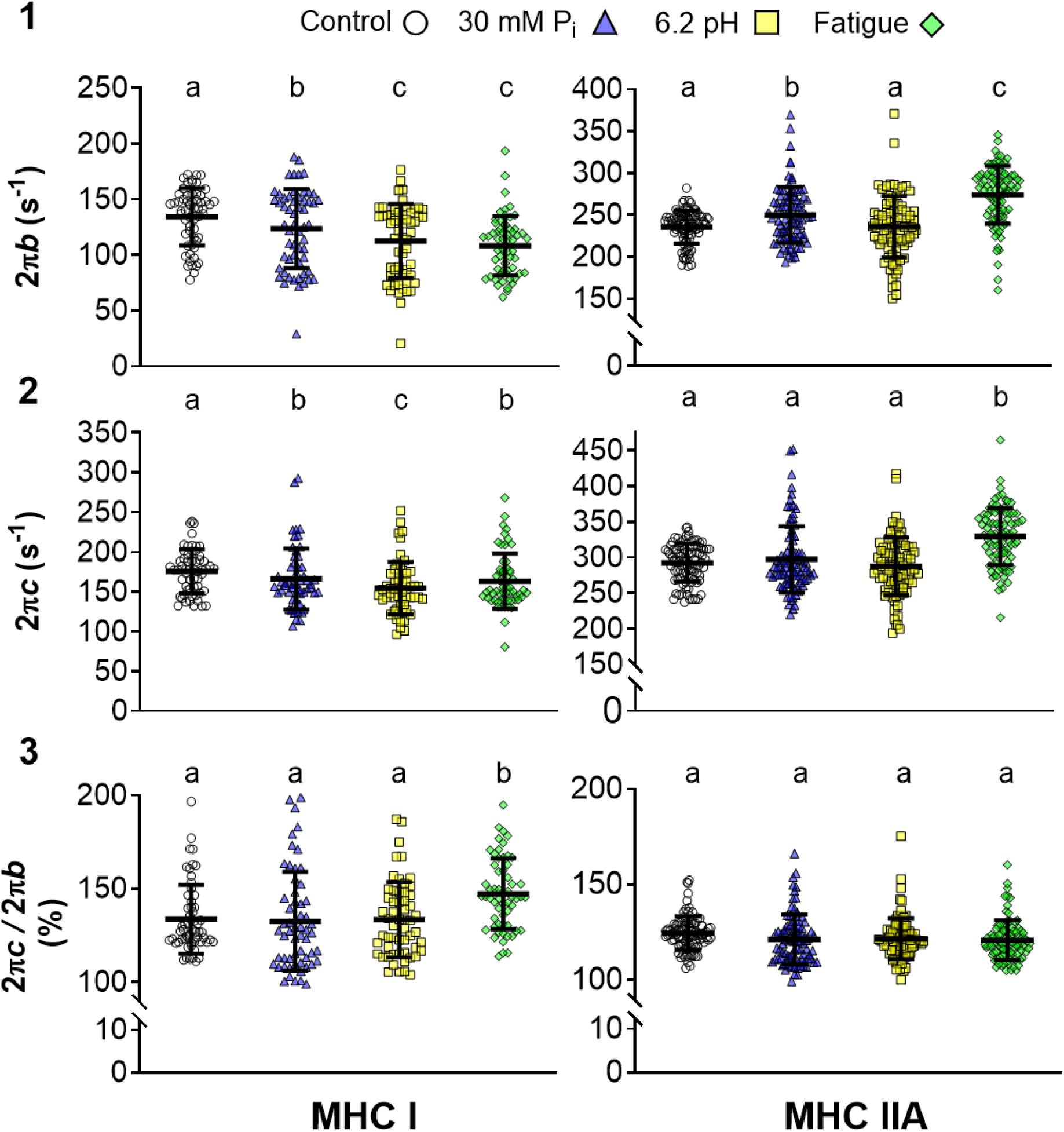
**Single-fiber maximum Ca^2+^-activated (pCa 4.5) sinusoidal analysis model kinetic parameters and ratios by MHC isoform for Control, high P_i_, low pH, and fatigue conditions at 37°C**. (1) 2π*b* is the rate of myosin transition between the weakly- and strongly-bound states, (2) 2π*c* is the rate of myosin detachment from actin. (3) 2π*c* / 2π*b* is the ratio of the kinetic rate constants of cross-bridge detachment, and cross-bridge recruitment. Data are presented as mean ± SD with individual fibers as data points. Letters above bars indicates significant (*p* < 0.05) pair-wise differences between conditions for MHC I or IIA.

In MHC IIA fibers, high P_i_ increased 2π*b* (*p* < 0.001) but not 2π*c* (*p* = 0.380). Low pH did not alter cross-bridge on- (*p* = 0.827) or off-kinetics (*p* = 0.270). In contrast, fatigue increased both 2π*b* (*p* < 0.001) and 2π*c* (*p* < 0.001), indicating that P_i_ and pH have an interdependent effect on cross-bridge kinetics, and an opposite response compared with the slowing of kinetics observed in MHC I fibers. These changes in cross-bridge kinetics did not alter their ratios (2π*c* / 2π*b*) (*p* = 0.796-0.952) except for fatigue in MHC I fibers, where this ratio increased (*p* < 0.001) (Figure 4.3), suggesting that the slowing of cross-bridge detachment rate exceeded the slowing of the rate of force production.

### Oscillatory Work Analysis

Oscillatory work was examined to understand how the six model parameters impact cross-bridge function when summed together. Oscillatory work was reduced from control in high P_i_ and low pH conditions for both MHC I and IIA fibers (Figure 5.1, Table 2), becoming negative in MHC IIA fibers. Reduced work was due to the decreased *B* and *C* discussed above and shown in Figure 2. The larger decrease in work in MHC IIA compared with MHC I fibers was likely caused by the increase in *C* / *B* in the individual high P_i_ and low pH conditions (Figure 3), as increasing the proportion of work absorbed will decrease the overall work produced. The frequency of peak work was reduced for pH but not P_i_ in MHC I fibers (Table 2), likely due to the greater slowing of cross-bridge kinetics with pH compared to P_i_ (Figure 4.1-4.2). In MHC IIA fibers, the frequency of peak work was reduced for both P_i_ and pH conditions, likely due to an increase in *C* / *B* (Figure 3.3) as cross-bridge kinetics were either faster or unchanged (Figure 4.1 and 4.2) and the 2π*c* / 2π*b* ratio was unchanged (Figure 4.3) compared to control.

**Figure 5.**
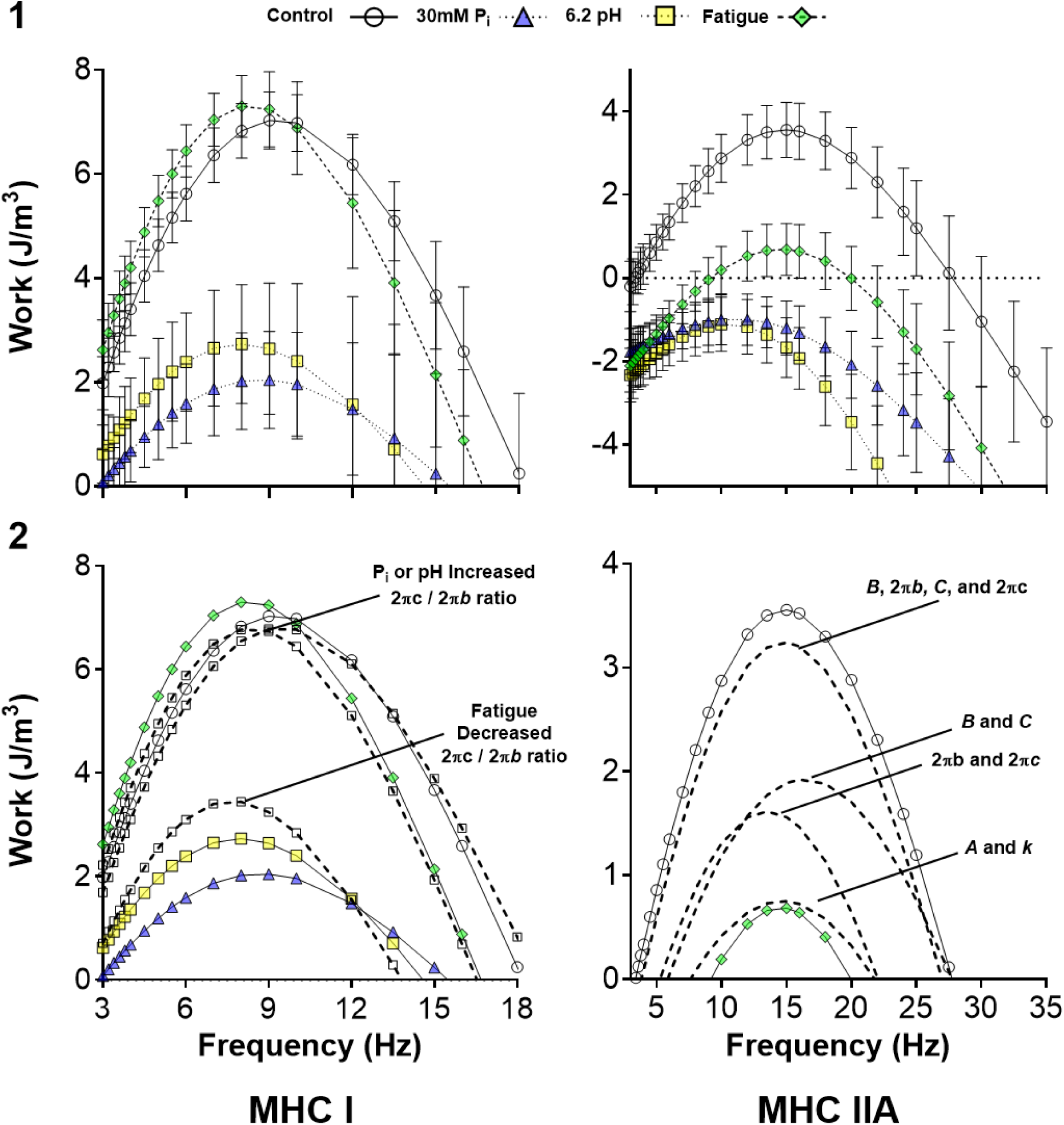
Single-fiber maximum Ca^2+^-activated (pCa 4.5) oscillatory work vs. frequency by MHC isoform for Control, high P_i_, low pH, and fatigue conditions at 37°C. (1): Oscillatory work for control, low pH, high P_i_, and fatigue by MHC isoform. (2): Models altering the 2π*c* / 2π*b* ratio (Left, MHC I) or the different model parameters (Right, MHC IIA) and their impact on work vs. frequency. Data are presented as mean ± SD. Models are compared to initial conditions of control, fatigue, high P_i_, and low pH for MHC I and control and fatigue for MHC IIA. Horizontal dotted line in upper right hand panel indicates zero work.

**Table 2.**
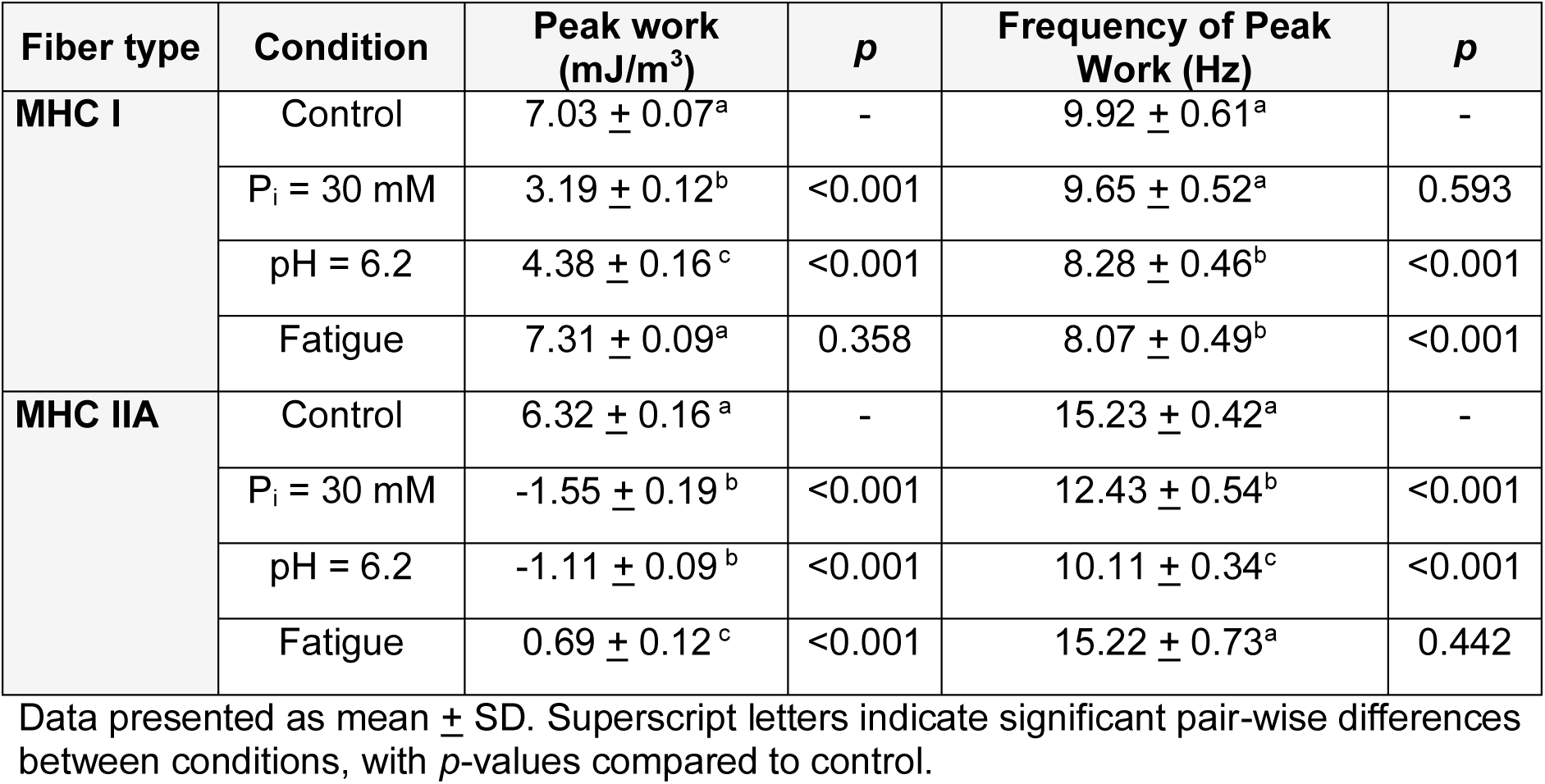
Summary of oscillatory work for MHC I and MHC IIA fibers.

| Fiber type | Condition | Peak work (mJ/m<sup>3</sup>) | *p* | Frequency of Peak Work (Hz) | *p* |
| --- | --- | --- | --- | --- | --- |
| **MHC I** | Control | 7.03 ± 0.07<sup>a</sup> | - | 9.92 ± 0.61<sup>a</sup> | - |
|  | P<sub>i</sub> = 30 mM | 3.19 ± 0.12<sup>b</sup> | <0.001 | 9.65 ± 0.52<sup>a</sup> | 0.593 |
|  | pH = 6.2 | 4.38 ± 0.16<sup>c</sup> | <0.001 | 8.28 ± 0.46<sup>b</sup> | <0.001 |
|  | Fatigue | 7.31 ± 0.09<sup>a</sup> | 0.358 | 8.07 ± 0.49<sup>b</sup> | <0.001 |
| **MHC IIA** | Control | 6.32 ± 0.16<sup>a</sup> | - | 15.23 ± 0.42<sup>a</sup> | - |
|  | P<sub>i</sub> = 30 mM | -1.55 ± 0.19<sup>b</sup> | <0.001 | 12.43 ± 0.54<sup>b</sup> | <0.001 |
|  | pH = 6.2 | -1.11 ± 0.09<sup>b</sup> | <0.001 | 10.11 ± 0.34<sup>c</sup> | <0.001 |
|  | Fatigue | 0.69 ± 0.12<sup>c</sup> | <0.001 | 15.22 ± 0.73<sup>a</sup> | 0.442 |

Data presented as mean ± SD. Superscript letters indicate significant pair-wise differences between conditions, with *p*-values compared to control.

Fatigue caused fiber-type specific effects such that in MHC I fibers a minimal change in oscillatory work, but a decreased frequency of max work, was noted, whereas in MHC IIA fibers the work was reduced with no change in frequency (Figure 5.1, Table 2). MHC I fibers under fatiguing conditions showed similar changes in most parameters compared to P_i_ and pH alone, except for cross-bridge kinetics, especially 2π*c* / 2π*b* which was increased with fatigue (147%) compared to P_i_ and pH alone (132-133%) (Figure 4.3). To determine the impact of 2π*c* / 2π*b* on work production, we created a hypothetical scenario where the 2π*c* fit parameter values were increased for P_i_ and pH alone such that their 2π*c* / 2π*b* values were increased to the fatigue value of 150% (Figure 4.3), while the remaining parameters (2π*b*, *B*, *C*, *A*, *k*) were left unchanged. By only increasing one parameter, 2π*c*, work curves for P_i_ and pH resembled the work curve for fatigue (Figure 5.2). Similarly, if the 2π*c* value for fatigue was reduced such that the 2π*c* / 2π*b* value was decreased to the approximate P_i_ and pH alone value of 130%, while the remaining parameters (2π*b*, *B*, *C*, *A*, *k*) were left unchanged, this work curve resembled the work curve for P_i_ and pH alone (Figure 5.2).

MHC IIA fibers under fatiguing conditions had reduced peak work compared to control, with the major changes to the 6-parameter model being a higher *k* and faster cross-bridge kinetics, 2π*b* and 2π*c*. To determine the impact of altering these parameters on oscillatory work production, the fit parameter values from control were substituted into the fatigue condition for *A* and *k* alone, 2π*b* and 2π*c* alone, *B* and *C* alone, or *B*, *C*, 2π*b*, and 2π*c* combined (Figure 5.2).

By only increasing *A* and *k*, work production was modestly increased, indicating changes to myofilament stiffness are not large contributors to the decrease in work in MHC IIA fibers. By altering 2π*b* and 2π*c* or *B* and *C* independently, work is shifted substantially higher relative to fatigue but does not completely explain the reduction in work with fatigue. While the mean values of *B, C,* and *B* / *C* were not statistically different between control and fatigue groups (Figure 2.1-2.1 and 3.3), the slight alterations to *B* / *C* from 107% to 105% in this model made fibers less work-absorbing and more work-producing. Thus, when altering *B* and *C* in conjunction with 2π*b* and 2π*c,* work production is returned almost to control conditions.

## DISCUSSION

We quantified cellular and molecular contractile function at 37°C in single human skeletal muscle fibers with the manipulation of P_i_ and pH independently and combined. Specific tension in both fiber types was reduced only under the fatigue condition (P_i_ = 30 mM, pH = 6.2) (Figure 1 and Table 3). Mechanisms leading to reductions in specific tension were fiber-type specific, with MHC I fibers displaying a reduced number of strongly-bound cross-bridges and/or stiffness; whereas MHC IIA fibers had greater myofilament and/or cross-bridge viscosity, leading to more work-absorption. Cross-bridge kinetics also responded differently by fiber-type, with MHC I fibers slowing in all three conditions (high P_i_, low pH, and fatigue) but MHC IIA fibers quickening with high P_i_ and fatigue. These alterations to molecular mechanics and kinetics led to reduced oscillatory work in both MHC I and IIA fibers when independently altering P_i_ or pH. But, with fatigue, oscillatory work was either returned to control for MHC I, or closer to control in MHC IIA. The mechanisms behind changes to work with fatigue were also fiber-type specific, with MHC I fibers increasing the ratio of the cross-bridge detachment rate to the rate of myosin force production, whereas MHC IIA fibers had alterations to work production and absorption processes (B- and C-processes). These findings highlight that (1) MHC I and IIA have a similar reduction in force in response to fatigue, but (2) the molecular mechanisms leading to reductions in force, and potential alterations to contractile velocities, are fiber-type specific at physiological temperature.

**Table 3.** Summary of independent and combined effects of reduced pH and increased P_i_ on cellular and molecular contractile function.

| Variable | 30 mM $P_i$ | | 6.2 pH | | Fatigue | |
| --- | --- | --- | --- | --- | --- | --- |
|  | I | IIA | I | IIA | I | IIA |
| **Specific tension (mN/mm<sup>2</sup>)** | ↔ | ↔ | ↔ | ↔ | ↓↓ | ↓↓ |
| **Strongly-bound cross-bridges (N/mm<sup>2</sup>)** | ↓↓ | ↓↓ | ↓↓ | ↓↓ | ↓↓↓ | ↔ |
| **Myofilament stiffness** | ↑ | ↑↑ | ↑ | ↑↑ | ↑ | ↓↓ |
| **$2\pi b$ (s<sup>-1</sup>)** | ↓ | ↑ | ↓↓ | ↔ | ↓ | ↑↑ |
| **$2\pi c$ (s<sup>-1</sup>)** | ↓ | ↔ | ↓↓ | ↔ | ↓ | ↑ |
| **Peak Work (J/m<sup>3</sup>)** | ↓↓↓ | ↓↓↓↓ | ↓↓ | ↓↓↓↓ | ↔ | ↓↓↓ |

↑, increase; ↔, no change, ↓, decrease; number of arrows signifies magnitude of response, with one arrow indicating 10-20% change, two arrows 20-50%, three arrows >50-100%, and four arrows >100%. Strongly-bound cross-bridges are derived from parameters $B$ and $C$ and myofilament stiffness from parameters $A$ and $k$ . $2\pi b$ is the rate of cross-bridge attachment and $2\pi c$ is the rate of cross-bridge detachment. Peak oscillatory work is derived from sinusoidal analysis at the lowest value of the viscous modulus.

Our results at 37°C show that cross-bridge kinetics responded to Pi and H^+^ differently by fiber type, with MHC I fibers slowing and MHC IIA becoming faster (Figure 4.1 and 4.2 and Table 3). Another widely used technique to examine isometric cross-bridge turnover kinetics is the rate of force development (Gordon *et al*., 2000), commonly called *k*_tr_, which incorporates both cross-bridge attachment and detachment rates (Brenner, 1988; Regnier & Homsher, 1998). Prior studies of fatigue in human MHC I and IIA fibers have shown reduced *k*_tr_ at 15°C (Sundberg *et al*., 2018) and decreased 2π*b* and 2π*c*, observed as increased myosin attachment time (*t*_on_, which equals (2π*c*)^-1^ (Palmer *et al*., 2013)), at 25°C (Foster *et al*., 2025). The opposite fatigue response in MHC IIA fibers at 37°C in the present study is likely due to temperature effects, as our previous work at 25°C (Foster *et al*., 2025) was performed in the same laboratory using the same equipment and in tissue from healthy older adults. In agreement with our fast-contracting MHC IIA results at 37°C, fatigued fast-contracting chicken pectoralis muscle (primarily MHC IIB, (Verdiglione & Cassandro, 2013)) had a faster rate of myosin detachment, using a mini-ensemble laser assay at 30°C (Woodward & Debold, 2018). This contrasts with their previous work using the same tissue and high temperature that showed fatigue slows unloading shortening velocity in the in vitro motility assay (Debold *et al*., 2011, 2012), suggesting a slower rate of myosin detachment, assuming a detachment-limited model of motility (Huxley, 1990). However, their recent work using fast-contracting rabbit psoas (primarily MHC IIX, (Tikunov *et al*., 2001)) using the laser trap assay at 25°C shows a slowed rate of detachment is likely due to an increased resistive load increasing the rate of P_i_ re-binding to myosin’s active site (Marang *et al*., 2025). This observation suggests that P_i_ alone will affect cross-bridge kinetics more under loaded conditions, such as in a mini-ensemble laser assay or an isometrically-contracting fiber (Reconditi *et al*., 2004), than in unloaded conditions, such as an in vitro motility assay. Thus, conditions that produce high resistive loads on the myosin, the mini-ensemble (Woodward & Debold, 2018) and an isometrically-contracting fiber, show an increased rate of myosin detachment with fatigue. In summary, our finding of slower cross-bridge kinetics in MHC I fibers at 37°C agrees well with previous human work (Sundberg *et al*., 2018; Foster *et al*., 2025). Therefore, assuming a greater resistive load on actomyosin in an AM.ADP state increases its affinity for P_i_ (Marang *et al*., 2025), our observation of faster cross-bridge kinetics with fatigue in MHC IIA fibers also agrees well with previous animal work (Woodward & Debold, 2018).

Previous studies have also found that cross-bridge kinetics respond to P_i_ or pH alone in a fiber-type specific manner (Millar & Homsher, 1992; Potma *et al*., 1995; Wahr *et al*., 1997; Tesi *et al*., 2000, 2002). Increasing P_i_ alone in slow-contracting fibers or myofibrils caused *k*_tr_ to be unchanged over the range 0-20 mM (Wahr *et al*., 1997; Tesi *et al*., 2000, 2002), but decreased at 30 mM (Wahr *et al*., 1997). Increasing P_i_ alone in fast-contracting fibers caused a very different response, with *k*_tr_ increasing at lower concentrations (Wahr *et al*., 1997; Tesi *et al*., 2000, 2002), and then not changing at concentrations > 10 mM (Wahr *et al*., 1997). The exponential rate of isometric force decline when caged P_i_ is released by laser flash photolysis, or *k*_Pi_, represents the effect that P_i_ binding has on strongly-bound cross-bridges (Dantzig *et al*., 1992). Slow-contracting fibers showed similar rates for *k*_Pi_ and *k*_tr_, while fast-contracting fibers showed higher rates for *k*_Pi_ compared to *k*_tr_ (Millar & Homsher, 1992), providing further evidence that P_i_ alone alters cross-bridge kinetics in a manner that depends on fiber type. Examining pH alone, the myosin ATPase rate slows in slow-contracting fibers and quickens in fast-contracting fibers contracting isometrically (Potma *et al*., 1995). In contrast, other experiments in fast-contracting animal fibers have shown that cross-bridge kinetics are unchanged with pH at 15°C (Metzger & Moss, 1990) and 32°C (Westerblad *et al*., 1997). Overall, these studies indicate that increased P_i_ or decreased pH alone show that cross-bridge kinetics decrease in slow-contracting fibers and increase or remain unchanged in fast-contracting fibers, in agreement with our current work at 37°C.

Although we have highlighted studies that generally agree with our current results, there are studies that report cross-bridge responses to metabolites that are different from those reported here, potentially due to differences in temperature, species, MHC isoform, or measurement technique. Temperature has a large effect on cross-bridge kinetics (Momb et al., Submitted companion paper) and their response to metabolites (Zhao & Kawai, 1994; Wang & Kawai, 2001; Sundberg *et al*., 2018), making comparison of results at different temperatures complicated. Cross-bridge kinetics and contractile velocities tend to slow as species become larger (Nyitrai *et al*., 2006; Pellegrino et al., 2003), suggesting that responses to metabolites in human fibers may be slower than in rabbit or mouse tissue. Indeed, the effect of increasing P_i_ in MHC I fibers at 37°C using sinusoidal analysis was variable in different species. Our current results in humans show 2π*b* decreases from 5 to 30 mM P_i_ (Figure 4.1), while rabbit soleus tissue increased 2π*b* between 0-30 mM P_i_ (Wang & Kawai, 2001). In addition to species differences, MHC isoforms have faster cross-bridge kinetics in the order MHC I < IIA < IIX < IIB (Galler *et al*., 2005) and can also show different responses to metabolites. For instance, the rate of myosin detachment increases with greater P_i_ in rabbit MHC IIA fibers, but remains unchanged with more P_i_ in MHC IIX and IIB fibers, despite the skeletal muscle fibers coming from the same species, rabbits (Galler *et al*., 2005). This isoform difference is especially important as numerous studies have used rabbit psoas tissue (primarily MHC IIX (Tikunov *et al*., 2001)), chicken pectoralis muscle (primarily MHC IIB (Verdiglione & Cassandro, 2013), or rat superficial vastus lateralis (primarily MHC IIB fibers (Metzger & Moss, 1987)) to examine the effects of metabolites on skeletal muscle cellular and molecular performance (Metzger & Moss, 1987, 1990; Cooke *et al*., 1988; Kawai & Halvorson, 1991; Millar & Homsher, 1992; Zhao & Kawai, 1994; Pate *et al*., 1995; Potma *et al*., 1995; Wahr *et al*., 1997; Tesi *et al*., 2000, 2002; Debold *et al*., 2008, 2011, 2012; Caremani *et al*., 2008; Woodward & Debold, 2018; Marang *et al*., 2025) (Kawai & Halvorson, 1991; Caremani *et al*., 2008).

In addition to species and MHC isoform effects, measurement technique may also play a role in metabolite responses. For instance, *k*_tr_ and sinusoidal analysis are quite different as the first uses a large amplitude slack-procedure to remove all strongly-bound myosin heads prior to allowing force development (Brenner, 1988) and the second uses oscillations small enough to probe cross-bridge kinetics without forcibly detaching the myosin heads (Kawai & Brandt, 1980; Palmer *et al*., 2007). These experimental differences can make interpretation difficult, especially when multiple steps in the cross-bridge cycle may change in response to metabolites or temperature. For instance, our fatigue results here at 37°C in human fibers show cross-bridge kinetics (2π*b* and 2π*c*) slow in MHC I fibers and become faster in MHC IIA fibers (Figure 4.1 and 4.2 and Table 3), while fatigue at 30°C in rat soleus and gastrocnemius show cross-bridge kinetics (*k*_tr_) in MHC I, IIA and IIX are unchanged compared to control (Nelson *et al*., 2014).

Similarly, our pH results show cross-bridge kinetics slow in MHC I fibers, whereas lowering pH at 15°C in rat soleus (primarily MHC I (Talmadge & Roy, 1993)) does not change cross-bridge kinetics (*k*_tr_) (Metzger & Moss, 1990). Whether these experimental results vary due to temperature, species, measurement technique, or another variable is not known at present.

Future work using a variety of techniques should focus on determining the effects of metabolites on cellular contractile function and myosin-actin cross-bridge kinetics in human skeletal muscle at *in vivo* temperature, as this should provide a clearer picture of the effects of fatigue and the ramifications for human health.

Single-fiber specific tension is the total force established by the number of strongly-bound cross-bridges, the force per cross-bridge and the stiffness of the myofilaments (Linari *et al*., 2004; Capitanio *et al*., 2006; Brenner *et al*., 2012). In this study, specific tension was not altered by manipulating P_i_ or pH independently, but was diminished 20-26% by their combination (Figure 1) due to a reduction in strongly-bound cross-bridges and/or their stiffness in MHC I fibers (Figure 3) and an increase in myofilament and/or cross-bridge viscosity in MHC IIA fibers (Figure 2.4). Reductions in force with fatigue are thought to be due to fewer force-generating cross-bridges and/or reductions in force per cross-bridge (Nelson *et al*., 2014; Foster *et al*., 2025), aligning well with the current findings in MHC I fibers. In contrast to MHC I fibers, however, MHC IIA fibers had an increase in myofilament and/or cross-bridge viscosity that resulted in reduced specific tension (Figure 2.4). These findings for MHC IIA are disparate from our prior work at 25°C, where the decrease in force in MHC IIA was due to a reduction in cross-bridge number (Foster *et al*., 2025). This result in MHC IIA fibers may be explained by faster cross-bridge kinetics (Figure 4.1-4.2), as previous work has identified an inverse relationship between cross-bridge lifetime and *k* (Wang *et al*., 2013; Palmer *et al*., 2013), such that faster cross-bridge kinetics increase myofilament viscosity, and thus, reduce force generation.

When altering P_i_ or pH independently, force production has been shown to markedly decrease at 15°C, whereas depression of force is only modest at 30°C or higher (Westerblad *et al*., 1997; Coupland *et al*., 2001; Debold *et al*., 2004; Knuth *et al*., 2006), aligning well with our results of no change at 37°C (Figure 1). Reductions in force at the cross-bridge level with elevated P_i_ have been proposed to occur due to early cross-bridge detachment (Marang *et al*., 2025), resulting in a decrease in the number of strongly-bound cross-bridges at the fiber level (Kawai & Halvorson, 1991; Caremani *et al*., 2008). However, P_i_ alone did not depress force in the current study, due to the counterbalancing effect of reduced cross-bridge number and/or stiffness with increased myofilament stiffness (Figures 1 and 2). Our findings that low pH did not lower force could be due to the rate of cross-bridge attachment being more sensitive to increased temperature than the rate of cross-bridge detachment (Wang & Kawai, 2001; Debold *et al*., 2004). Thereby, the purported mechanism of decreased force with pH -slowing of the forward rate constant (Metzger & Moss, 1990; Woodward & Debold, 2018; Jarvis *et al*., 2018) - may be mitigated at higher temperatures. Notably, the combined effect of both high P_i_ and low pH led to a decrease in specific tension in both fiber types and thus is the likely explanation for muscle fatigue *in vivo*.

To understand how changes to the underlying cross-bridge mechanics and kinetics combine to alter contractile function, we analyzed the performance of molecular motors by measuring oscillatory work at 37°C. When manipulating P_i_ or pH independently in MHC I and IIA fibers, reduced or negative work occurred, with no change in specific tension (Figure 5)., This was a result of reduced *B* and *C* in MHC I fibers, but an increased *C* / *B* ratio in MHC IIA fibers (Figures 2 and 3). While not a direct comparison, depression of oscillatory work occurs without reduced force production in *Drosophila* with mutations to flightin (Henkin *et al*., 2004), indicating performance of molecular motors can change absent of alterations to force. Our findings indicate that the ability to generate work may be compromised when altering P_i_ or pH alone, even if cellular level force is not changing substantially. Notably, oscillatory work during fatigue conditions in MHC I and IIA fibers was similar or closer to control conditions, indicating an interacting effect of these metabolites. In MHC I fibers, this occurs because as the oscillatory frequency of the C-process increases, the amount of work absorption at the lower oscillatory frequencies (where positive work output occurs) decreases. Thus, altering 2π*c* / 2π*b* with fatigue in MHC I fibers causes the fatigue response in these fibers to more closely resemble control than P_i_ or pH alone. In MHC IIA fibers, by modeling how transitioning 2π*b* and 2π*c, B* and *C*, or a combination of all four back to control levels, we found the greatest impact to work production via the combination of all four (Figure 5.4). While *B, C*, and their ratio were statistically different (Figure 3), the subtle change in *C* / B from 107% in fatigue to 105% in control created a shift from work absorption (C-process) to work production (B-process). Thus, while a primary difference between fatigue and control is an increase in myofilament viscosity, peak work with fatigue is reduced primarily due to alterations in work production and work absorption. These findings further quantify the interacting effects of P_i_ and pH at 37°C, with fiber-type specificity.

Importantly, MHC I fibers displayed a general fatigue-resistance regarding oscillatory work when high P_i_ and low pH were combined, highlighting important fiber-type specific differences to *in vivo* metabolite accumulation.

Our data bring to light new evidence of the effects of fatigue on human skeletal muscle single fiber contractile function at a physiological temperature (37°C). Specifically, molecular-level contractile function is different between slow- and fast-contracting fibers when altering P_i_ or pH independently or in combination. The fatigue properties of MHC I and IIA fibers is a crucial topic as these MHC isoforms make up a large proportion of human skeletal muscle (Murach *et al*., 2019). Thus, the proportion of each in an individual may influence overall muscle performance during fatigue (Thorstensson & Karlsson, 1976; Bagley *et al*., 2017). Our findings highlight that although both fiber types exhibit reductions in force production with fatiguing conditions, MHC I fibers may be fatigue resistant during oscillatory work production, whereas MHC IIA fibers are not. Because the mechanisms underlying the response to fatiguing conditions are fiber-type specific at 37°C, this indicates a need to examine force-velocity-power profiles at physiological temperature. While reductions in peak power occur in both fiber types by 40-55% in human single fibers (Sundberg *et al*., 2018), and by ∼60% in rat single fibers (Nelson *et al*., 2014), these studies were conducted between 15-30°C and our results suggest these findings may not translate to *in vivo* temperature in humans. If feasible, future work should aim to test single fiber function, or other molecular preparations, close to or at physiological temperature for comparison to parallel studies *in vivo*.

## ADDITIONAL INFORMATION

### Data availability statement

All data are represented within this manuscript.

### Author contributions

B.A.M and M.S.M. conceived and designed the research; B.A.M and S.R.C. acquired human skeletal muscle tissue; B.A.M. performed experiments; B.A.M. analyzed data; B.A.M, J.A.K, S.R.C., and M.S.M interpreted results of research; B.A.M., J.A.K and M.S.M prepared figures; B.A.M. and M.S.M. drafted manuscript; All authors contributed to the critical revision of the manuscript and approved the final version of the manuscript submitted for publication.

## Acknowledgements

We thank the volunteers who dedicated their valuable time to these studies and the Center for Human Health and Performance in the Institute for Applied Life Sciences at the University of Massachusetts, Amherst. We also thank Professor Edward “Ned” Debold for helpful discussions and comments on the manuscript.

## Author’s present address

Department of Kinesiology 30 Eastman Lane Totman Building, Room 140E University of Massachusetts Amherst, MA 01003 USA

